# Emergence and Dynamics of Usutu and West Nile Viruses in the Netherlands, 2016-2022

**DOI:** 10.1101/2024.12.16.628479

**Authors:** Emmanuelle Münger, Nnomzie Atama, Jurrian van Irsel, Rody Blom, Louie Krol, Tijs J van den Berg, Marieta Braks, Ankje de Vries, Anne van der Linden, Irina Chestakova, Marjan Boter, Felicity D Chandler, Robert Kohl, David F Nieuwenhuijse, Mathilde Uiterwijk, Ron A M Fouchier, Hein Sprong, Andrea Gröne, Constantianus J M Koenraadt, Maarten Schrama, Chantal B E M Reusken, Arjan Stroo, Judith M A van den Brand, Henk P van der Jeugd, Bas B Oude Munnink, Reina S Sikkema, Marion P G Koopmans

## Abstract

**Background:** Mosquito-borne arboviruses, including Usutu virus (USUV) and West Nile virus (WNV), are emerging threats in Europe, with changes in climate, land use, and increased travel and trade influencing their dynamics. Understanding the emergence and establishment of these viruses in new regions is critical for informing targeted mitigation of drivers of emergence and enhancing public and wildlife health preparedness.

**Methods:** Seven years (2016-2022) of interconnected studies were conducted in the Netherlands. Live birds were sampled by volunteer bird ringers, dead birds were referred for sampling by citizen-scientists and zoological institutions, and mosquitoes were trapped. Samples were tested for USUV and WNV using RT-PCR and screened for *Orthoflavivirus* antibodies using protein microarray and neutralization assays. Sequencing and phylogenetic analyses were performed.

**Findings:** USUV was first detected in the Netherlands in 2016, caused large outbreaks in birds until 2018, and resurged in 2022. One dominant, enzootic lineage, co-circulated alongside several limited introductions of a second lineage. A localized WNV lineage 2 outbreak occurred in live birds and mosquitoes in 2020, with another positive bird in 2022 and serological evidence of continued circulation.

**Interpretation:** We provide the first comprehensive, multi-year documentation of two emerging arboviruses in the Netherlands. USUV was enzootically maintained over seven years and was associated with substantial bird mortality. WNV is in an early stage of establishment with no bird mortality observed. Our integrated wildlife sampling was crucial in detecting a human WNV outbreak, bringing this emerging threat to public health attention.

**Funding:** ZonMw, Dutch Research Council (NWO), European Union, Government of the Netherlands

## Introduction

Mosquito-borne arboviruses can cause severe disease in humans, domestic animals, and wildlife. These viruses are maintained in infection cycles involving mosquitoes as vectors, and vertebrates as reservoir hosts. Ongoing global changes in climate, land use, travel, and trade will likely increasingly affect the interactions of arboviruses, their vectors, and their hosts, with a rise in viral prevalence expected in temperate regions^1^. Therefore, arboviral diseases are listed as climate-sensitive priority diseases^1,2^. In recent years, mosquito-borne arboviruses have expanded to and within Europe^3^ as a result of i) geographical expansion and establishment of arboviruses in resident bird and native mosquito populations, ii) outbreaks following the importation of viruses by travellers to regions with established invasive *Aedes* mosquito species, and iii) effects of climate warming, changing precipitation patterns, and other environmental shifts on habitats suitability for arbovirus maintenance. Understanding the processes of emergence and establishment of arboviruses in new regions is crucial to allow development of risk-based early-warning detection and prediction systems, particularly in regions with occasional, self-limiting arbovirus activity on the verge of local establishment.

Usutu virus (USUV) and West Nile virus (WNV) are examples of regionally emerging arboviruses. Both are mosquito-borne *Orthoflaviviruses* and belong to the Japanese encephalitis virus serocomplex. In Europe, *Culex pipiens* mosquitoes are their main vectors. While birds serve as reservoirs for these viruses, some species are also highly susceptible to disease. Eurasian Blackbirds (*Turdus merula*) and captive Great Grey Owls (*Strix nebulosa*) have been particularly affected by USUV outbreaks in Europe^4^. WNV incursion into the Americas caused large-scale declines in bird populations, with members of the crow family (*Corvidae*) and other passerines particularly affected^5^. In contrast, in Europe, as in Africa and Asia, WNV has not been reported to cause high mortality in birds.

Humans (*Homo sapiens*) and other mammals are considered dead-end hosts of USUV and WNV; they generally do not develop sufficient viremia to infect mosquitoes and do not contribute to the transmission cycle. Most human infections with USUV and WNV are asymptomatic, however WNV can cause severe disease, with ∼20% of infected individuals developing febrile illness, and up to 1% neurological symptoms^6^. Symptomatic USUV cases are rare, but are characterized by fever, jaundice, rash, or neurological complications (reviewed by Cadar & Simonin^7^).

USUV was first detected in Europe in 2001, when it was identified as the causative agent of mass bird mortality in Austria^8^. USUV has since spread to most European countries, with multiple lineages present across the continent^9^. Human WNV cases were first detected in Europe in 1958 through serosurveys in Albania^10^. Early outbreaks in Europe were primarily associated with WNV lineage 1^11^. In 2004, WNV lineage 2 was detected in Hungary and rapidly became dominant on the continent, causing large outbreaks in humans and animals^11^. WNV is currently enzootic in several European countries^9^.

Aside from South of France^12^, there were no reports of either USUV or WNV in Western Europe before the first USUV outbreak in Austria in 2001^8^. Since then, USUV has emerged in several Western European countries^9^ and recently also reached Sweden^13^, Denmark^14^, and the United Kingdom^15^. Emergence of USUV preceded emergence of WNV in Austria^16^, Germany^17^, Switzerland^18^ and, as we will describe in this study, the Netherlands. Given the similarity of WNV and USUV transmission cycles, vector species, and host ranges, as well as antigenic cross-reactivity between the two viruses, USUV circulation may indicate environmental suitability for WNV, and the circulation of one virus could influence that of the other^19^.

While global change is expected to influence the suitability of temperate regions for mosquito-borne arboviruses, key aspects of these pathogens’ dynamics in newly susceptible regions –including host range breadth, potential for enzootic persistence, overwintering mechanisms, or environmental drivers of viral dynamics- remain poorly understood. Hence, in-depth studies in a wide range of wildlife hosts and vectors on the moving front of these pathogens are essential to address these gaps. As arboviruses can circulate in wildlife well before human or veterinary disease is observed, such studies may also serve as early-warning surveillance system.

Here, we present the results of seven years of research on mosquito-borne viruses in birds and mosquitoes across the Netherlands, between 2016 and 2022. Aiming to detect and understand the introduction and spread of emerging arboviruses, we established a nationwide integrated sampling framework in a unique collaboration of ecologists, ornithologists, entomologists, veterinarians, virologists, and citizens. A multi-tiered sampling scheme was designed to collect samples from free-ranging live and dead birds, birds deceased in captivity, and mosquitoes. We screened these samples for arboviruses using molecular and serological methods. We describe the emergence of USUV and WNV, their avian host range and their epidemiology in the Netherlands.

## Research in Context

### Evidence before this study

We searched PubMed for articles including the terms “detection”, “emergence”, “cases”, “circulation”, “prevalence” or “presence” in combination with either “arbovirus”, “Usutu virus”, “USUV”, “West Nile virus”, or “WNV” and a country of Western Europe: “Austria”, “Belgium”, “France”, “Germany”, “Liechtenstein”, “Luxembourg”, “Monaco”, “Netherlands”, “Switzerland”. This search yielded 128 relevant articles. No articles were found regarding Liechtenstein and Monaco. Articles highlighted the recent emergence of Usutu virus (USUV) in all other countries, and of West Nile virus (WNV) in Austria, France, Germany, the Netherlands, and Switzerland, illustrating the ongoing geographic expansion of these viruses. There was very limited longitudinal data following initial outbreak reports, mainly from Germany and Austria. Many studies focused on either molecular or serological investigation, in one specific type of host (e.g., wild birds, captive birds, equids, human blood donors) or in mosquitoes.

### Added value of this study

This study provides a comprehensive, seven-year longitudinal analysis of data from live and dead birds as well as mosquitoes, documenting the emergence and subsequent spread of USUV and WNV in avian hosts and vectors in the Netherlands with exceptional detail. It leverages a nationwide sampling framework that integrates public engagement, ecology, ornithology, entomology, and virology, and combines molecular and serological methods. Through its inclusive approach, it provides data on involvement and impact of the target viruses in multiple host species, a unique feature allowing comparative analysis across host species and viruses. Our findings reveal the enzootic nature of USUV, associated with mortality in bird populations, and indicate the likely early establishment of WNV in the country – a presence that routine surveillance had failed to detect, underscoring the unique reach of our integrated approach. They further contribute to defining the host range of both arboviruses and provide an excellent basis for studying key drivers of expansion in a region previously considered free from USUV and WNV.

### Implications of all the available evidence

The combined evidence from this study and existing literature highlights the expanding geographic range, increasing incidence, and establishment of arboviruses in local bird populations and native mosquitoes in Europe. This expansion is likely underpinned by climate change and other environmental changes. These findings underscore the need for preparedness for climate-sensitive emerging infectious diseases, to mitigate a possible greater toll of arboviral disease on human and other animal populations in the future. Our findings, along with others’, underscore the importance of arbovirus studies in wildlife hosts and vectors for detecting emerging arboviral threats to public health, as demonstrated by the identification of human WNV cases following our findings in birds and mosquitoes. These wildlife study efforts could be optimized for application in other settings. Integrating arbovirus findings with climate, host, and landcover covariates could improve our understanding of the conditions driving mosquito-borne virus dynamics and, ultimately, enhance predictive models for outbreaks, inform targeted intervention strategies, and develop robust outbreak risk indicators.

## Methods

We generated data about circulation of arboviruses through the studies described below (**Figure 1**). The studies were spatially and temporally connected, with samples processed using the same protocols.

**Figure 1:**
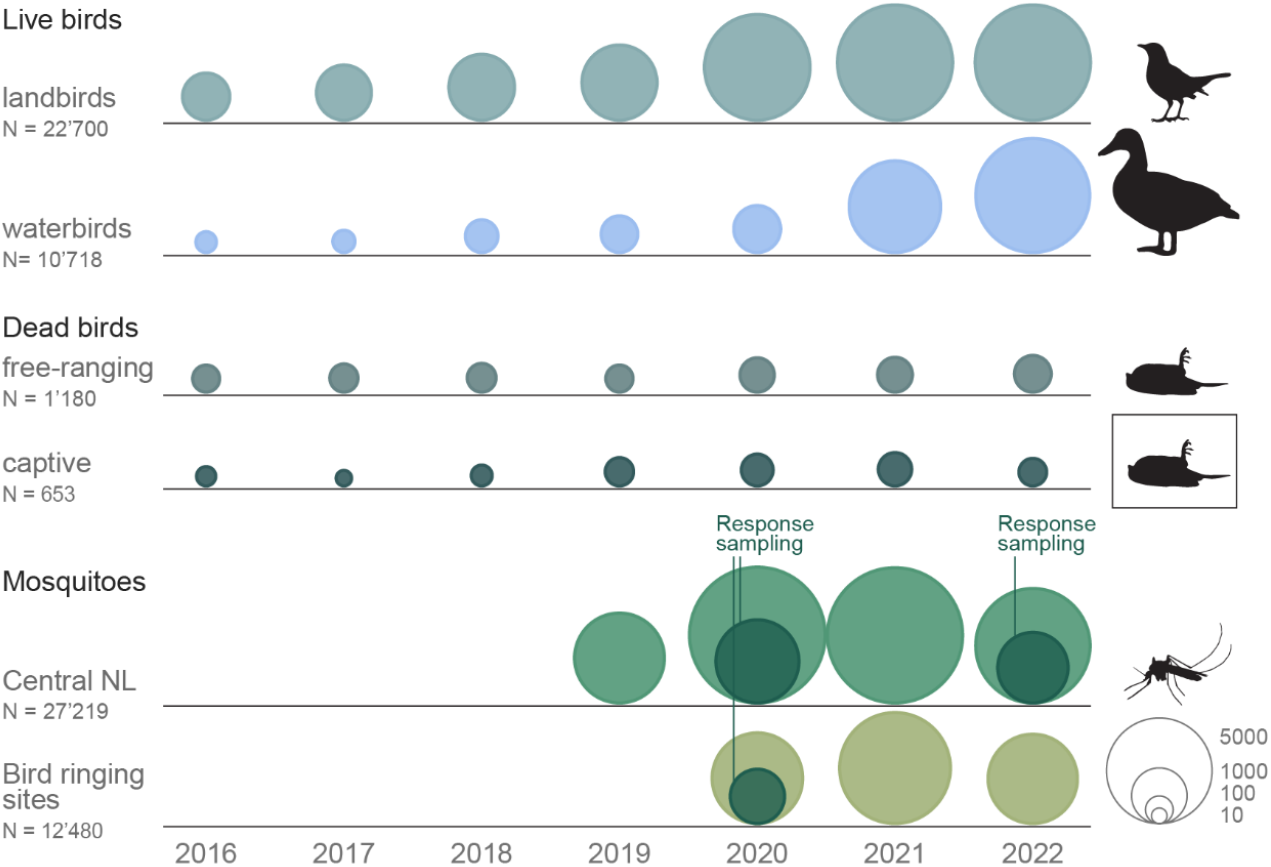
Overview of the different sampling schemes in birds and mosquitoes through time. The total number of samples tested per study are indicated (N). Circle sizes are proportional to the number of samples tested each year.

Additional details on the methods described hereafter are provided in **appendix 1**.

### Citizen-science based sampling of live free-ranging birds

Live free-ranging birds were captured and sampled by trained volunteer ornithologists across the Netherlands from March 2016 to December 2022 (**Figure S5**), under ethical permits. Birds were ringed, weighed and measured. Optimally, throat and cloacal swabs, along with a blood sample (whole blood and/or dried blood spots), were collected. Additionally, from 2020, feather samples were collected. Recapture and resampling of ringed birds occurs frequently during the breeding season, and the numbers presented in this study reflect the number of sampling events of individual birds, rather than the absolute number of individual birds. From 2020, arbovirus testing was added for birds originally sampled for avian influenza virus monitoring to explore utility of a combined approach; this significantly increased numbers of waterbirds tested. For analyses, bird species were categorized as “waterbirds” or “landbirds” based on orders.

### Citizen-science and zoological institution-based referral of dead birds for pathology

Dead free-ranging birds were reported across the Netherlands through a citizen science system^20^ from March 2016 to December 2022 (**Figure S6**). Dead captive birds were submitted by zoos and veterinarians. Dead birds were necropsied, and samples were collected. Initially, dead birds were submitted for USUV and WNV diagnostics if necropsy indicated USUV as a possible cause of death; from 2019, all submitted birds were tested.

### Mosquito collection

Mosquitoes were trapped weekly at bird ringing sites throughout the Netherlands between July and October 2020–2022 using fermenting molasses-baited traps, as previously described^21^. In addition, weekly trapping was carried out in the central Netherlands between May and October 2019–2022, using CO^2^ baited light traps. Mosquitoes were identified to species level and pooled to a maximum of 10 individuals by species (complex/group) and trapping event. Following WNV detections in Utrecht (2020) and Hollands Kroon (North Holland, 2022), sampling was intensified around detection locations until the end of the mosquito season (October).

### USUV and WNV diagnostics

Samples from dead birds (with brain tissue as preferred sample), live birds (throat and cloacal swabs), and mosquito pools were screened using real-time PCRs (RT-PCR) for the presence of USUV and WNV.

### Antibody detection in serum from live free-ranging birds

Bird sera were screened for USUV and WNV antibodies in a protein microarray (PMA) approach as previously described^22^. Sera with a signal above background and sufficient sample volume were tested further using Virus Neutralization Tests (VNT) and Focus Reduction Neutralization Tests (FRNT) as previously described^22,23^. A serum sample was considered positive for WNV- or USUV-neutralizing antibodies when reactivity on PMA was confirmed by FRNT or VNT, with a titre at least 4-fold higher for one virus. Samples with less than a 4-fold difference were classified as *Orthoflavivirus* positive.

### Viral whole genome sequencing and phylogenetic analysis

USUV and WNV RT-PCR positive wildlife samples with CT values below 32 were submitted to whole genome sequencing using an amplicon-based Oxford Nanopore approach, as previously described^24^. Additionally, USUV positive human blood donations from the Netherlands, 2018^25^ were followed up by sequencing. Near full length USUV and WNV lineage 2 genomes were retrieved from GenBank^26^, aligned with newly generated sequences, and maximum likelihood phylogenetic analyses were conducted.

### Spatial analysis

For details on clustering and aggregation of sampling locations, see appendix 1.

## Results

### Molecular detections in birds

Between March 2016 and December 2022, samples from 22’700 live landbirds, 10’718 live waterbirds, 1’180 dead free-ranging birds and 653 dead captive birds were tested for USUV and WNV. USUV was detected in 156 live landbirds (0.7%), 3 live waterbirds (0.03%), 243 dead free-ranging birds (20.6%) and 41 dead captive birds (6.3%). USUV was detected in 29 species of free-ranging birds, primarily from the order Passeriformes, with 156 detections in live birds and 233 detections in dead birds across 22 species within this order (**appendix 2**). Eurasian Blackbirds had the highest USUV prevalence among live birds (n=106/4’415, 2.4%) and the highest number of deaths associated with USUV infection (n=208). A selection of species with strong evidence of USUV infection is presented in **Figure 2**. While USUV was incidentally detected in live Anseriformes and Charadriiformes, prevalence in these orders, intensively sampled after 2020, was low (n=1/7’819 and n=1/2’048 respectively). In dead captive birds, USUV was detected in 17 species, with Great Grey Owls (*Strix nebulosa*, order Strigiformes) showing particularly high positivity (n=20/22).

**Figure 2:**
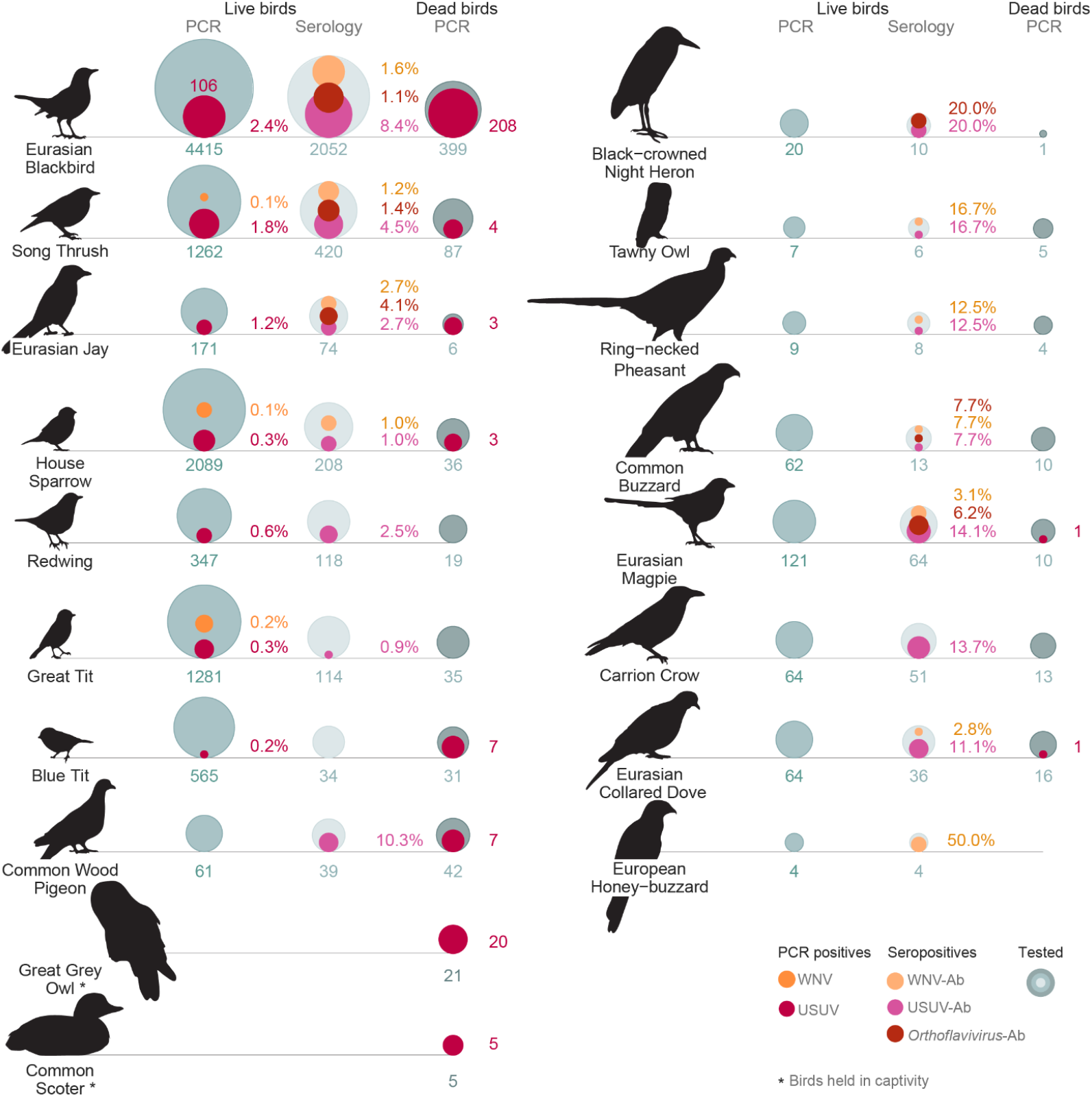
Selection of bird species, both free-ranging and captive, with evidence of Usutu virus (USUV) or West Nile virus (WNV) infection and/or exposure, the Netherlands, 2016-2022. Species are selected based on: (i) more than 1 USUV detection and prevalence greater than 0.25% in live birds, or (ii) more than 2 USUV detections in dead birds, or (iii) more than 1 USUV and/or WNV seropositive bird and seroprevalence greater than 10%. Ordering of species from top to bottom and left to right follows: USUV prevalence in live and/or dead birds and *Orthoflavivirus* seroprevalence. To allow visual differentiation of small values, circle size is proportional to the square root of numbers tested or positive. Total number tested, prevalence, and seroprevalence for live birds, and number of cases for dead birds are labelled for each species; a complete overview for all species tested is provided in **appendix 2**.

WNV was detected in 7 live landbirds (one of which tested positive twice, with a 5-days interval) from 5 species within the order Passeriformes, as well as in 1 live waterbird, a Grey Heron (*Ardea cinerea*, order Pelecaniformes - **appendix 2**). WNV was not detected in any dead bird. USUV and WNV detections overlapped in 5 bird species.

### Serological detections in live free-ranging birds

For a subset of 4’176 live landbirds (18.4%), a serum sample was collected. 240 landbirds tested positive for USUV-neutralizing antibodies (5.7%), 53 for WNV-neutralizing antibodies (1.3%) and 40 tested positive for both USUV- and WNV-neutralizing antibodies (1%), without the possibility to discriminate between the two. USUV antibodies were detected in 26 species, and WNV antibodies in 14 species, with overlapping detections in 10 species. Eurasian Blackbirds had an USUV seroprevalence of 8.4% (n=172/2052). Among the 10 species with strongest evidence for exposure to USUV and/or WNV (more than one bird positive for antibodies and seroprevalence greater than 10%), 8 had no or incidental molecular detections (**Figure 2**, right column). It is noteworthy that these species were tested in relatively small numbers, both in live and dead birds.

### Molecular detections in mosquitoes

Between 2019 and 2022, 39’699 mosquitoes from 20 species, primarily *Cx. pipiens/torrentium* (88.39%), were tested in 6’480 pools. USUV was detected in 28 mosquito pools (27 *Cx. pipiens/torrentium*, and 1 *Anopheles maculipennis* s.l), and WNV in 6 pools, all *Cx. pipiens/torrentium* (**Table S1**).

### Spatiotemporal patterns of Usutu virus and West Nile virus detection

USUV was first detected in the Netherlands in spring 2016^27^ and was associated with major die-offs of free- ranging and captive birds in summer 2016. Since then, USUV was detected annually in live and dead, free- ranging and captive birds. Mosquito collection, implemented since 2019, also yielded USUV-positive pools every summer (**Figure 3A, Figure 4**). Numbers of samples tested and positive samples per research project and year are presented in **Table S2**.

**Figure 3:**
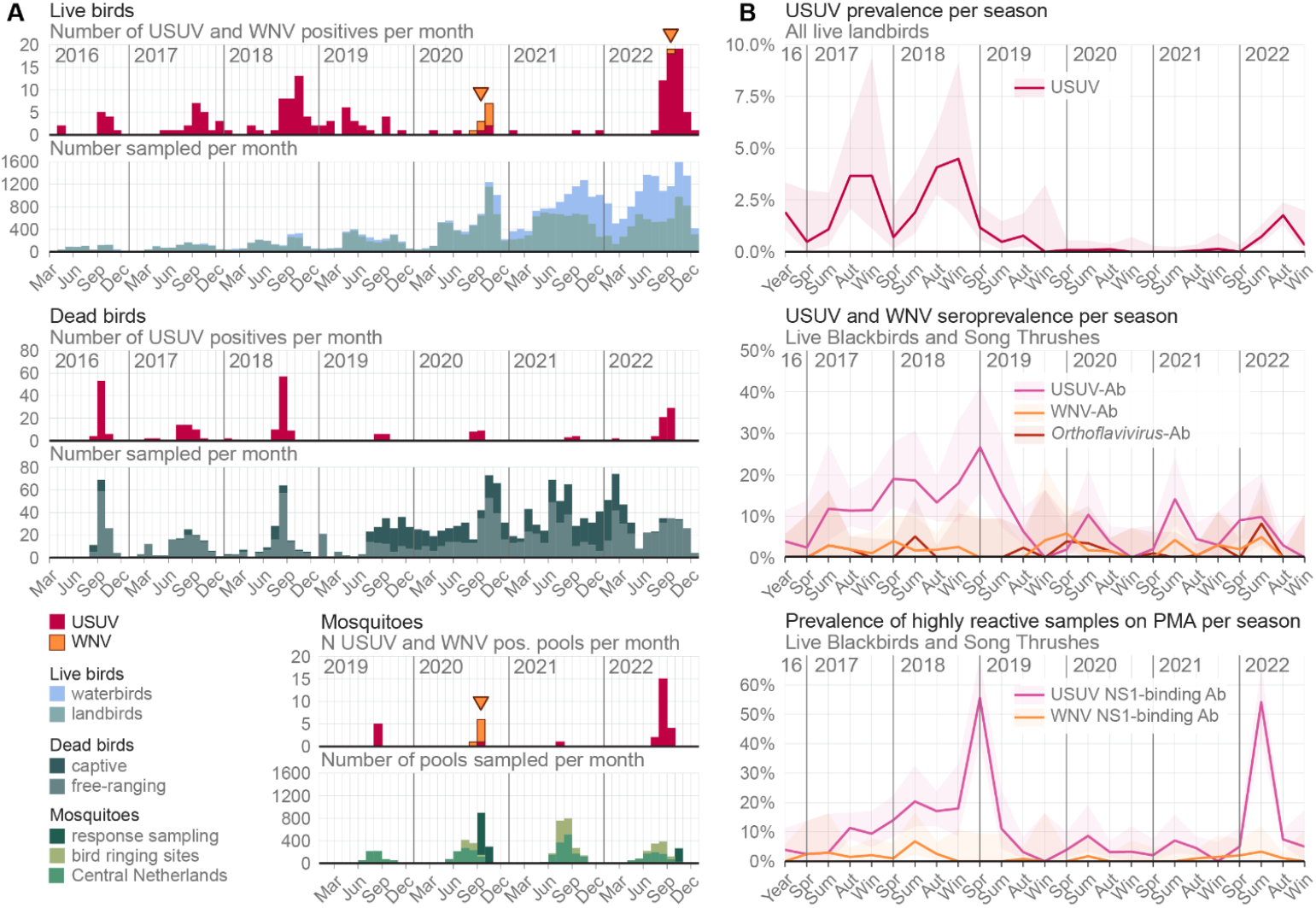
Temporal dynamics of Usutu virus (USUV) and West Nile virus (WNV) in the Netherlands, 2016-2022. **(A)** Temporal patterns of USUV and WNV incidence in the different hosts and vectors studied. Top panel: survey results in live birds; middle panel: dead birds; lower panel: mosquitoes. Each histogram represents the number of cases per month, with colour-coded bars indicating the virus type or host/project type. Orange arrows point to periods of WNV detections. **(B)** Prevalence and seroprevalence of USUV and WNV in live landbirds of the Netherlands across seasons. Top panel: prevalence of USUV in live landbirds, expressed as percentages of RT-PCR positive cases out of the total tested by RT-PCR; middle panel: seroprevalence for USUV and WNV in live Eurasian Blackbirds and Song Thrushes, expressed as percentages of neutralization assay confirmed seropositives out of the total tested on the protein microarray (PMA); lower panel: percentage of live Eurasian Blackbirds and Song Thrushes tested on USUV and WNV PMA with signal greater than 30’000 relative fluorescence units. Prevalence and seroprevalence estimates are shown as solid lines, shaded ribbons represent the 95% CI based on the Agresti-Coull method.

**Figure 4:**
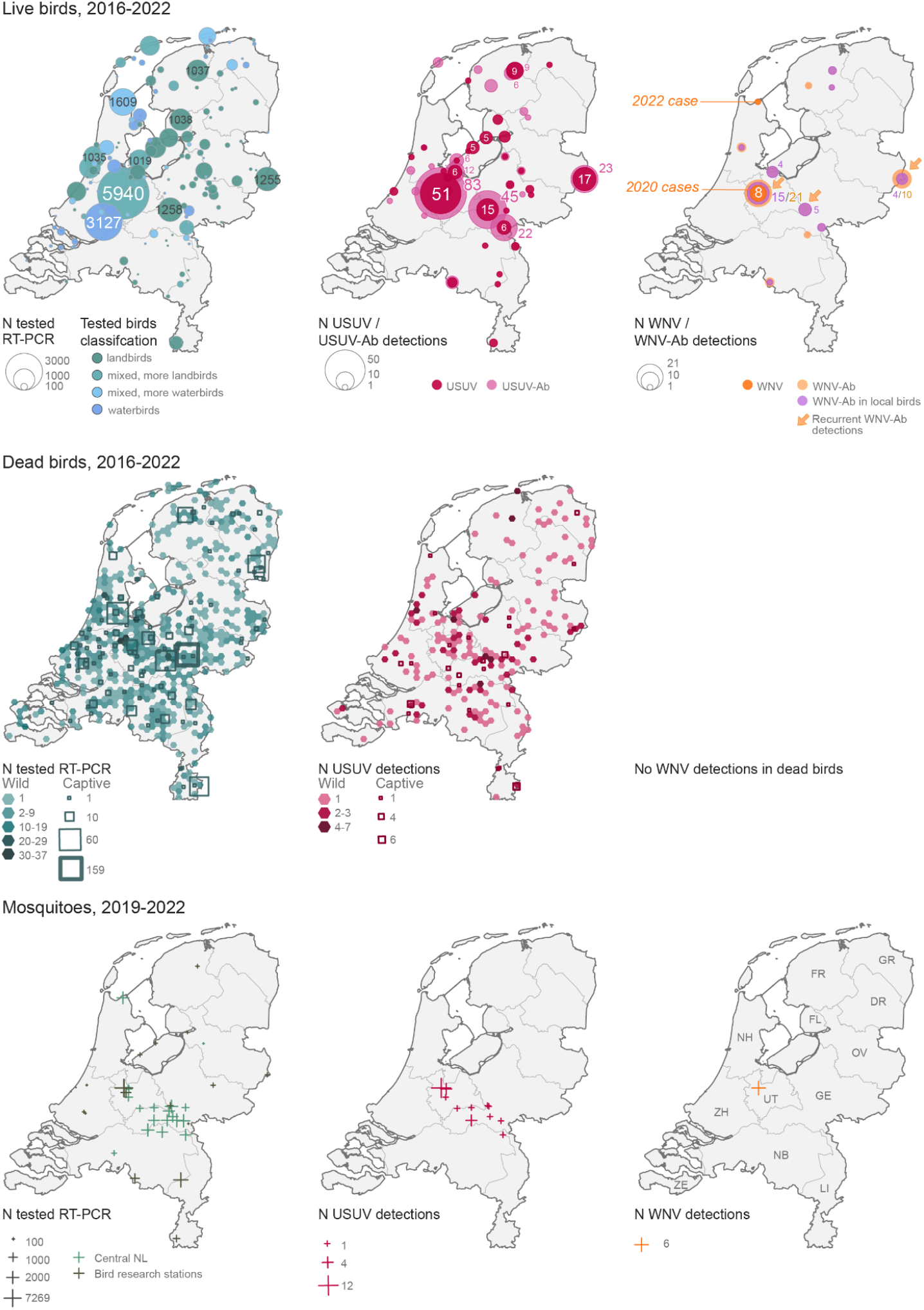
Geographical distribution of sampling and Usutu virus (USUV) and West Nile virus (WNV) detections in the Netherlands, 2016-2022. Upper row: live birds (2016- 2022); middle row: dead birds (2016- 2022); lower row: mosquitoes (2019-2022). Left maps show the number of individuals tested by RT-PCR, centre maps show USUV and USUV-neutralizing antibodies detections (“USUV-Ab”, live birds only) and right maps show WNV and WNV-neutralizing antibodies detections (“WNV-Ab”, live birds only); arrows indicate locations with WNV-Ab detections over multiple years. Different symbols are used for the different types of host/survey and sized proportionally to numbers. The colour scheme distinguishes between hosts sampled, virus and virus-neutralizing antibodies. Provincial borders are shown and are labelled with two-letter abbreviations in the lower right map. Source administrative boundaries: CBS, Kadaster, “CBSGebiedsindelingen2022” (https://service.pdok.nl/cbs/gebiedsindelingen/atom/v1_0/index.xml).

The highest numbers of USUV detections in birds occurred from 2016 to 2018, peaking in 2018 with cases in 39 live landbirds (2.5% of 1574 tested) and 72 dead free-ranging birds (60.5% of 119 tested). Initially, USUV-positive birds were predominantly detected in the southern and central regions of the Netherlands, shifting to central and northern regions in 2018 (**Figure S1**). After this, the number of cases decreased. 2022 saw a resurgence in cases mainly in the southern and central regions, with USUV detections in 55 live landbirds (0.9% of 6148 tested) and 54 dead free-ranging birds (18.7% of 289 tested). Despite continued mosquito sampling throughout the country, USUV was detected only in mosquitoes in the central region, possibly related to high numbers of mosquitoes collected from these locations and an overall low prevalence in mosquitoes. USUV outbreaks were seasonal, with most detections in birds occurring between July and November. However, detections also occurred during the colder months, with 14 live and 4 dead free-ranging birds testing positive for USUV from December to February.

Eurasian Blackbirds and Song Thrushes (*Turdus philomelos*) were the species most abundantly and consistently tested by serology throughout the study period. Consequently, the temporal serological pattern is primarily informed by these species. Increased USUV seroprevalence was observed between 2017 and 2019, peaking at 26,7% in Eurasian Blackbirds and Song Thrushes in spring 2019, then dropping by autumn 2019 (**Figure 3B)**. In 2016, serum testing was very limited; in 2017, USUV-neutralizing antibodies were predominantly detected in the southern and central regions, shifting to the central and northern regions in 2018 and 2019 (**Figure S2, Figure S3**). Serum sampling remained scarce in the southern and northeastern most regions throughout the study period. The prevalence of highly reactive sera on the USUV PMA mirrored the overall seroprevalence trend, with the notable exceptions of two peaks, in spring 2019 and summer 2022 (55% and 54% highly reactive, respectively).

In August 2020, a Common Whitethroat (*Curruca communis*) and 2 *Cx. pipiens/torrentium* pools sampled in Haarzuilens (Utrecht) tested positive for WNV lineage 2^28^. Additional WNV detections in September and October brought the total to 8 detections in live birds and 6 detections in mosquito pools during this local outbreak. WNV was not detected in 2021 but was again detected in 2022 in a single live Grey Heron, in North Holland (**Figure 3A, Figure 4**).

WNV-neutralizing antibodies were detected annually in live free-ranging birds since 2017, at low prevalence. At the country level, no increase in WNV seroprevalence was observed following the 2020 detections (**Figure 3B**). Evidence of exposure to WNV was found in a total of 53 birds, in Utrecht (the site of the 2020 WNV detections) as well as in 10 additional locations. Based on species and capture histories, 35 of these seropositive birds were identified as local (tested between 2016 and 2022). In Utrecht, WNV- neutralizing antibodies were identified in 1 to 6 birds annually between 2017 and 2022, with seroprevalence ranging from 0.7% to 2.9% and 15 of 21 seropositive birds identified as local (tested between 2018 and 2022). Recurrent detection of WNV-neutralizing antibodies was also observed at two other locations, in the provinces of Overijssel and Gelderland (**Figure S4**). At the first location, 3 of 10 seropositive birds (tested in 2017, 2019, and 2022) and all 4 seropositive birds at the second location (tested in 2016, 2020, and 2022) were identified as local, suggesting local WNV circulation.

### Phylogenetic analyses

In total, 247 near full-length USUV genome sequences were generated through the studies described, derived from 35 live birds, 183 dead free-ranging birds, 23 dead captive birds, and 6 mosquito pools. In addition, 3 near full-length and 2 partial USUV genomes sequences were generated from positive human blood donors. 133 new USUV genome sequences are released in this study, 119 sequences have been released before^24,29^.

Our phylogenetic analysis shows that USUV lineages Africa 3 and Europe 3 co-circulated in the Netherlands, with Africa 3 predominantly detected (**Figure 5 A**). Lineage Africa 3, with clades primarily composed of sequences from the Netherlands, shows a phylogenetic structure suggesting enzootic establishment. Re-emergence of closely related strains was observed year after year between 2016 and 2019. From 2020 onwards, a specific sub-cluster of USUV Africa 3 was most frequently detected. For lineage Europe 3, the structure of the phylogenetic tree, with several small clades of sequences from the Netherlands separated by sequences primarily from Germany (2011 to 2016), suggests multiple introductions of this lineage into the country. The same lineages were detected in live and dead free-ranging birds, dead captive birds, mosquitoes, and humans.

**Figure 5:**
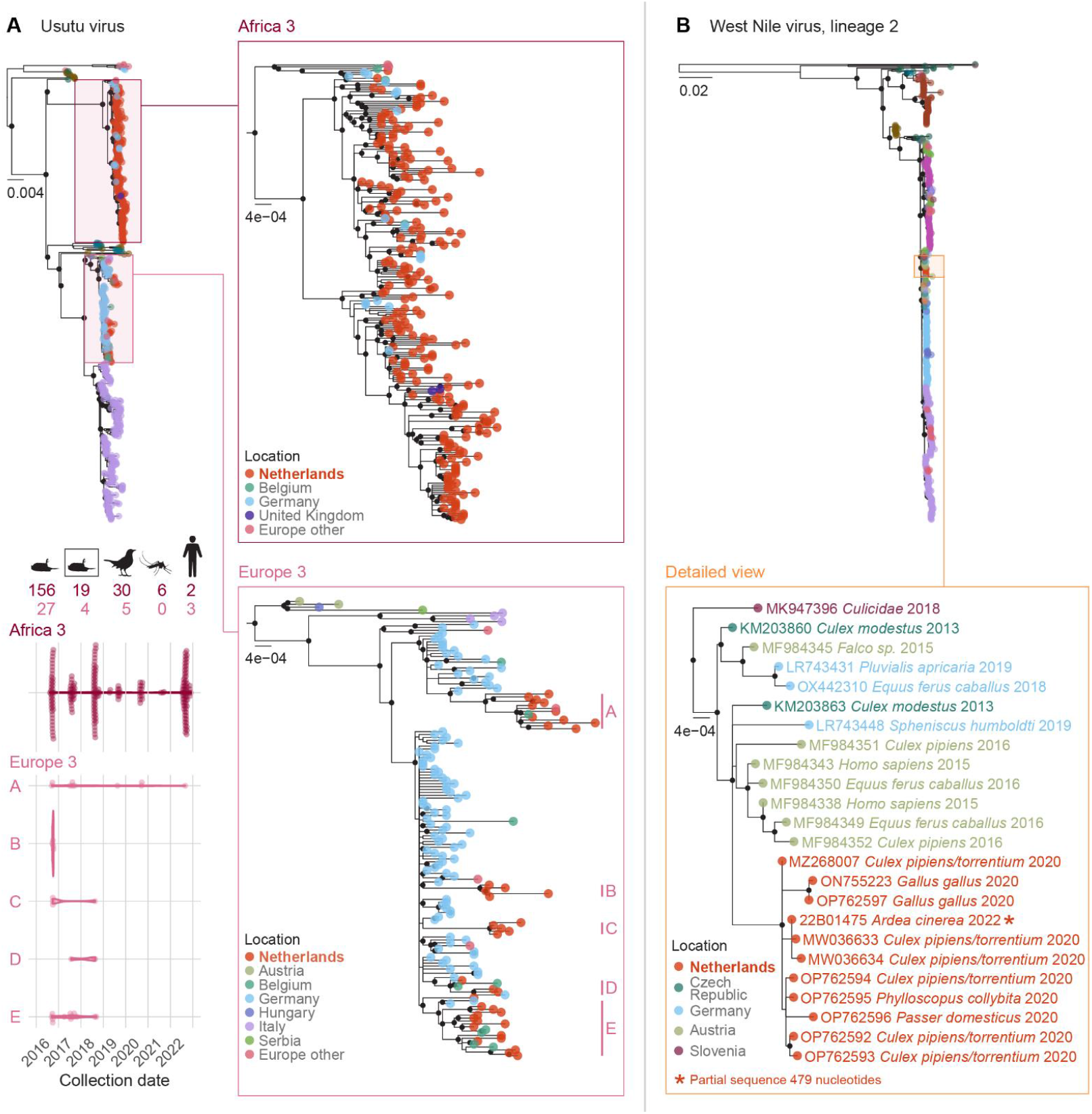
Phylogenetic analyses of Usutu virus (USUV) and West Nile virus (WNV) lineage 2 genome sequences from hosts in the Netherlands, 2016-2022 (A) Maximum likelihood phylogeny of USUV complete coding sequences, with expanded views of lineages Africa 3 and Europe 3. Clades predominantly composed of sequences from the Netherlands are labelled A-E. In the lower left, the temporal distribution of sampling dates for isolates from the Netherlands is shown by clade; the number of sequences derived from (1) dead free-ranging birds, (2) dead captive birds, (3) live birds, (4) mosquitoes, and (5) humans is indicated. Colour intensity reflects lineage classification, with Africa 3 in dark pink and Europe 3 in light pink. (B) Maximum likelihood phylogeny of WNV complete coding sequences, including one partial WNV genome from the Netherlands (2022), with an expanded view of the clade containing sequences from the Netherlands. Phylogenetic tree tip colours indicate the geographic origin of each sequence, nodes with bootstrap support >90 are labelled with a dot, and the unit of scale is substitutions per site.

Seven near full-length WNV lineage 2 genome sequences were obtained from 2 live birds and 6 mosquito pools from the 2020 outbreak in Utrecht; all sequences from this outbreak were closely related (**Figure 5 B**; 2 additional sequences from a chicken associated to this outbreak are included). Sequences from the Netherlands were more distantly genetically related to sequences from Germany (2019), Austria (2015- 2016) and the Czech Republic (2013). A partial genome sequence (479 nucleotides within the envelope protein coding region) was retrieved from the 2022 WNV-positive Grey Heron; it was identical to the corresponding region of sequences from the 2020 Netherlands outbreak.

## Discussion

Over the last decades, the geographic range of arboviruses in Europe has expanded and these viruses have caused an increasing number of outbreaks in humans and in animals. This emphasizes the need for a better, systemic understanding of their ecology and for the development of effective arbovirus monitoring approaches to inform public health and environmental strategies. Through spatially and temporally connected studies of arbovirus circulation in birds and mosquitoes, we show that USUV and WNV have emerged and established in the Netherlands within the last eight years. We describe their temporal dynamics and spatial spread and identify a broad range of host species.

Sampling free-ranging live birds, initiated in March 2016, provided early warning of the circulation of USUV in the Netherlands^27^. Five months later, the first bird deaths attributed to USUV foreshadowed mass mortality events amongst free-ranging and captive birds. Screening human blood donations in summer 2018, a period of intense USUV circulation in wildlife, led to the detection of human USUV infections^25^. We here show that these human infections were caused by lineages Africa 3 and Europe 3, which circulated simultaneously in birds in the Netherlands. Because USUV infection in birds often led to death, monitoring the virus in citizen-reported dead birds was especially valuable. Compared to monitoring the virus in live birds, testing of far fewer dead bird allowed for the detection and tracking of USUV’s spatiotemporal dynamics. Additionally, high viral loads in dead birds’ tissues increased viral genome sequencing success, making dead birds particularly valuable for genomic monitoring of USUV.

Combining molecular and serological studies in live birds with molecular studies in dead birds allowed us to comprehensively cover species that are involved in the ecology of USUV as reservoir and/or susceptible host. The strongest evidence of infection and associated mortality was observed in Eurasian Blackbirds, and high numbers of detections were noted in Great Grey Owls deceased in captivity, consistent with observations from other countries^4,19^. Our findings also add to the growing evidence of infection and exposure in a broader range of birds, including species of Corvidae, Columbidae, and raptors (Accipitridae and Strigidae). While USUV caused mass mortality in Eurasian Blackbirds, the frequent detection of USUV and USUV-neutralizing antibodies in living individuals without visible symptoms of disease suggests that this species can also carry the virus while remaining fit. Eurasian Blackbirds are the most common breeding birds in the Netherlands, and research in the USUV- and WNV- affected area of Northern Italy has shown a feeding preference of *Cx. pipiens* for this species^30^. They may therefore play a role in sustaining viral transmission to mosquitoes, despite the high mortality. However, the wide range of bird species with evidence of USUV infection or exposure underscores the need to investigate their role within the reservoir community, which may be essential for understanding USUV dynamics. Herons, crows, and raptors, with wider ranges of movement that cover extensive areas and longer lifespans than Eurasian Blackbirds, may play distinct roles in the transmission and spread of USUV.

USUV seroprevalence sequentially followed molecular prevalence, increasing from summer 2017 and peaking in spring 2019. Seroprevalence waned rapidly, declining by autumn 2019 and remaining at lower levels; this pattern may facilitate the recurrence of outbreaks at multi-year intervals. Peaks in prevalence of highly reactive sera on the PMA, seen in 2020 and 2022, that were unmatched by similar levels of USUV and WNV neutralizing antibodies, are notable and deserve further investigation. Perfect congruence between neutralization assays and NS1 binding is not necessarily expected, as these methods target antibodies recognizing different epitopes, which may persist in birds for varying durations. Studies on long-term kinetics of antibody responses to *Orthoflavivirus* infections in birds would aid interpretation of these findings. However, given the high antigenic cross-reactivity among *Orthoflaviviruses* antibodies, this may also suggest the circulation of another related virus.

The persistent presence of USUV in the Netherlands over 7 years, characterized by the dominant circulation of lineage Africa 3 (which is not known to circulate in Southern Europe) and the annual emergence of related strains strongly suggests enzootic circulation and overwintering of the virus in the country or broader Western European region. Our earlier phylodynamic analyses indicated that USUV had been circulating in the Netherlands or neighbouring regions years before it was detected in the Netherlands^24^. The resurgence of strains most closely related to those from the 2016-2018 transmission period, resulting in a new surge in cases in 2022, further supports enzootic maintenance and silent circulation preceding larger outbreaks. While USUV dynamics suggest it can overwinter locally, which may also apply for WNV, the mechanisms enabling persistence of these viruses through winter remain unclear.

USUV was recently detected in diapausing *Cx. pipiens*/*torrentium* in the Netherlands (*Koenraadt et al*., *under review*), and infected diapausing mosquitoes are considered the primary overwintering pathway for USUV and WNV^31,32^. Our studies detected high rates of USUV infections in birds in late autumn (after reduced mosquito activity), as well as several infections in winter, consistent with reports in outdoor aviaries in Germany^33^. This suggests long-term virus persistence in avian hosts. Persistent arboviral infections, observed with WNV in experimentally inoculated birds^34^ may also serve as an overwintering mechanism in temperate regions^34,35^. Additionally, bird-to-bird transmission in winter roosts^36^ and winter-active mosquitoes might contribute to the overwintering of USUV and WNV.

USUV dynamics in neighbouring Germany suggest a link to outbreaks in the Netherlands. USUV was first detected in Southwest Germany in 2011^37^. Similar to the Netherlands, bird cases in Germany increased in 2016, with the virus spreading to new areas, including regions bordering the Netherlands^38^. While lineage Europe 3 was predominant in Germany until 2018^38^, lineage Africa 3 became the main circulating lineage in 2019^38^. USUV lineage Europe 3 and Africa 3 have also been described in Belgium^39^ and lineage Africa 3 was recently detected in the United Kingdom^15^.

WNV was detected for the first time in the Netherlands in 2020, in Utrecht, in live free-ranging birds and mosquitoes^28^. Sampling live birds and mosquitoes proved crucial, as no dead birds with evidence of WNV infection have been found in the Netherlands to date. Following these detections, awareness was raised amongst health professionals, and retrospective analyses of cases of neurological disease of unknown aetiology were undertaken. This resulted in the identification of 8 symptomatic human cases in the country that year, 6 with West Nile virus neuroinvasive disease and 2 with West Nile fever^40,41^. In accordance with European regulations, screening of blood donations for WNV was initiated in the region of the index patients and adjacent regions. All blood donations tested negative for WNV^41^. Given the small proportion of human WNV infections that develop into neuroinvasive disease, it is likely that a larger epizootic outbreak occurred locally in 2020, that could have gone unnoticed without our studies in birds and mosquitoes. In 2022, WNV was again detected in a heron, with a (partial) genomic sequence closely resembling viruses from the 2020 outbreak, while WNV-neutralizing antibodies were repeatedly detected in local birds in Utrecht and two additional locations. These observations, along with detection of seroconversions in sentinel chickens in 2021 and 2022 in Utrecht and Gelderland^42^, indicate ongoing local circulation of WNV at low levels in different regions of the Netherlands.

As WNV detections in wildlife preceded detections in humans, this demonstrates that studies in birds and mosquitoes have potential for early warning surveillance, although unlike in North America, this was not detected based on bird mortality^43^. In Austria, increased USUV activity in birds was reflected in increased numbers of positive human blood donations, and close genetic relationship of USUV and WNV sequences was observed in human and birds populations^44^. The approaches used in ecological surveillance and the information these efforts can provide may thus differ based on factors specific to each region, such as patterns of arbovirus circulation, impact of arboviruses on wildlife and human health, and regional public and veterinary health priorities.

In conclusion, we describe the emergence and establishment of two mosquito-borne arboviruses, USUV and WNV, in a previously non-endemic country. This phenomenon is likely driven by changes in local environmental conditions, becoming favourable for arbovirus establishment in locally present (non-introduced) mosquito species, larger outbreaks in wild birds, and occasional zoonotic transmission. The intensity of arbovirus activity in a particular season is likely influenced by a combination of factors, including avian host abundance, variations in host immunity, and favourability of climatic conditions to mosquito and virus populations. Integrating arbovirus findings, including our USUV and WNV datasets, with ecological and climatic variables can refine predictive outbreak models, inform targeted interventions, and support identification of robust risk indicators. Methods such as ecological niche modelling, transmission models, and phylodynamic analyses can further improve our understanding of periods and areas favourable for the sustained circulation of USUV, WNV and related *Orthoflaviviruses*. Early detection of arboviruses in mosquitoes and birds can ensure timely implementation of prevention and control measures to protect human health. These include targeted vector control, public communication on personal protection measures, blood donations screening, raising awareness among health professionals, and the inclusion of the specific arbovirus in differential diagnosis of encephalitis cases.

## Contributors

**EM**: Conceptualization, Data Curation, Formal Analysis, Investigation, Methodology, Visualization, Writing – Original Draft; **NA, AvdL, IC, MBoter, FC, RK, DFN**: Investigation, Data Curation, Methodology (All studies, virology); **JvI, TJvdB**: Investigation, Data Curation, Methodology (Live birds studies); **RB, LK**: Investigation, Data Curation, Methodology (Mosquitoes at bird ringing sites); **MBraks, AdV, MU**: Investigation, Data Curation, Methodology (Mosquitoes in Central Netherlands); **RAMF, HPvdJ**: Conceptualization, Supervision (Live birds studies); **AG, JMAvdB**: Conceptualization, Supervision (Dead birds studies); **CJMK, MS**: Conceptualization, Supervision (Mosquitoes at bird ringing sites); **HS, AS**: Conceptualization, Supervision (Mosquitoes in Central Netherlands); **CBEMR**: Conceptualization, Supervision (Birds studies, virology); **BBOM**: Conceptualization, Supervision (All studies, Pathogen genomics), Writing – Original Draft; **RSS, MPGK**: Conceptualization, Supervision, Funding Acquisition (Overall project), Project Administration, Writing – Original Draft; **All authors**: Writing – Review & Editing

## Data availability

The complete datasets on live and dead free-ranging birds, birds deceased in captivity, and mosquitoes analysed in this study are available in the BioStudies database (http://www.ebi.ac.uk/biostudies) under accession numbers [accession numbers] and can be accessed via [Pathogens Portal Netherlands, currently in advanced development]. Supplementary tables regarding numbers of individuals tested and positive per bird and mosquito species are provided in the appendix (mosquitoes: appendix 1, Table S2; birds: appendix 2). Viral genome sequences and raw reads generated in this study have been deposited on The European Nucleotide Archive (ENA) under accession numbers [accession numbers].

## Acknowledgments

We sincerely thank the numerous volunteer bird ringers and citizen field assistants for collecting samples from live birds, mosquitoes, and associated metadata, as well as the general public for reporting wild birds’ mortality. We extend our thanks to zoos and veterinarians who submitted birds that died in captivity. We are grateful to Oanh Vuong, Sanne Thewessen, Mikaela Suehely Cicilia, and Seren Altundag at the Erasmus MC Rotterdam for their contributions to sample processing as well as to Tess van de Voorde, Jolien Morren and Natasja van Nijen at Vogeltrekstation NIOO-KNAW, who managed training, logistics, and communication for bird ringers. Lastly, we thank the staff of the Dutch Wildlife Health Centre and the Veterinary Pathology Diagnostic Centre in Utrecht, including Ruby Wagensveld-van den Dikkenberg, for processing the dead birds, as well as Timon ten Berge, Hanna Hesselink, and Natashja Ennen-Buijs for administration and data management related to wild bird mortality reporting.

This work was supported by the Eco-Alert project funded by ZonMw with project number 522001004; the research program One Health PACT (NWA.1160.18.210) which is (partly) financed by the Dutch Research Council (NWO); the European Union’s Horizon 2020 research and innovation program under grant agreement No. 874735 (VEO); The Dutch Ministries of (i) Agriculture, Fisheries, Food Security and Nature (ii) Health, Welfare and Sport (National Institute for Public Health and the Environment).

During the preparation of this work, the authors used ChatGPT for language refinement and text streamlining suggestions. The authors reviewed and edited the content as needed and take full responsibility for the final publication.

