## Appendix 1 for "Emergence and Dynamics of Usutu and West Nile Viruses in the Netherlands, 2016-2022"

### Supplementary Appendix 1 for

#### Emergence and Dynamics of Usutu and West Nile Viruses in the Netherlands, 2016-2022

Emmanuelle Münger, Nnomzie Atama, Jurrian van Irsel, Rody Blom, Louie Krol, Tijs J van den Berg, Marieta Braks, Ankje de Vries, Anne van der Linden, Irina Chestakova, Marjan Boter, Felicity D Chandler, Robert Kohl, David F Nieuwenhuijse, Mathilde Uiterwijk, Ron A M Fouchier, Hein Sprong, Andrea Gröne, Constantianus J M Koenraadt, Maarten Schrama, Chantal B E M Reusken, Arjan Stroo, Judith M A van den Brand, Henk P van der Jeugd, Bas B Oude Munnink, Reina S Sikkema, Marion P G Koopmans

**The PDF file includes:**

**Other Supplementary Materials for this manuscript include:**

Appendix 2: Breakdown of individuals sampled, viral detections, and antibody detections by bird order and species

### Methods

#### Animal Sampling, Pre-Diagnostic Processing and Analyses

Sampling of live free-ranging birds was conducted under ethical permits AVD801002015342 and AVD80100202114410 issued to NIOO-KNAW. For analyses, bird species from the orders Anseriformes, Charadriiformes, Ciconiiformes, Gruiformes, Pelecaniformes, Podicipediformes, and Suliformes were categorized as “waterbirds”, and other species as “landbirds”. For live landbirds and waterbirds, viral diagnostics were performed by RT-PCR on throat and cloacal swabs, with unconfirmed detections occasionally verified using a second sample type. Most waterbirds were tested only using molecular methods. Based on species and capture histories, birds with evidence of West-Nile virus exposure were classified as local, non-local, or unknow. Non-migratory, resident species were always considered local. Migrant species that are present in the Netherlands only in either summer or winter were always considered non-local. Blackbirds, which both breed and winter in the Netherlands were considered local if sampled outside the migration and winter season, or when additional captures of the same individual indicated that the bird was part of the local resident breeding population. In other cases, birds were categorized as unknown. Reported dead free-ranging birds were selected for necropsy at the Dutch Wildlife Health Centre (DWHC) based on freshness of the carcasses and likelihood of disease-related death. Between 2016 and 2018, following initial detections of USUV related mortality, dead free-ranging birds were preferentially selected for post-mortem investigation at locations where USUV activity had not been identified before. Dead captive birds were submitted for necropsy at the Veterinary Pathology Diagnostic Centre (VPDC). A detailed description of necropsies is provided in Giglia et al[1]. Testing for USUV and WNV diagnostics in dead birds was done by RT-PCR of brain tissues; if brain tissue was unavailable, diagnostics were occasionally performed on alternative tissues.

In the central region of the Netherlands, mosquitoes were trapped using $CO_2$ baited CDC miniature light traps (John. W. Hock Company, Gainesville, USA). At intensified sampling locations following WNV detections, CDC Gravid Trap Model 1712 (JW Hock) for collection of gravid females and mosquito aspirators were additionally used.

For nucleic acid extraction from live bird swabs, 600 µL of the viral transport medium (VTM) was used as input material. Tissue samples from dead birds were homogenized in tissue lysis buffer (MagNA Pure DNA Tissue Lysis Buffer) using the Fastprep bead beater (MP Biomedicals, shaking at 4.0 m/s for 10 s). 60 µL of the supernatant was mixed with 540 µL of VTM or Dulbecco's Modified Eagle Medium (DMEM) prior to nucleic acid extraction. Total nucleic acid (NA) was extracted using the MagNA Pure 96 Instrument and MagNA Pure 96 and DNA and Viral NA Large Volume Kit (Roche LifeScience, Basel, Switzerland) according to the manufacturer’s instructions. Mosquito pools where homogenized in 1 ml medium (DMEM, $NaHCO_3$, Hepes-buffered saline, penicillin/streptomycin, amphotericin) with one 1/4″ ceramic sphere (MP Biomedicals, Solon OH, USA) using the FastPrep-24 5G bead beater (5 m/s for 30 s). Homogenates were cleared by centrifugation. 60 µl of homogenate supernatant was added to 90 µl MagNA Pure 96 External Lysis Buffer (Roche), and nucleic acids were extracted using Ampure beads (Beckman Coulter Life Sciences, Indianapolis, IN, USA) and a DynaMag 96 side magnet (Invitrogen/Thermo Fisher, Waltham, MA, USA).

Samples from dead birds (with brain tissue as preferred sample), live birds (throat and cloacal swabs) and pools of mosquitoes trapped at bird ringing sites were screened using real-time PCRs (RT-PCR) for the presence of USUV[2] and WNV[3], positive results were confirmed by a second USUV[4] or WNV[5] RT-PCR targeting a different region of the genome. Phocine distemper virus was added as internal NA extraction control[6]. Pools of mosquitoes trapped in the central Netherlands were screened using RT-PCR for the presence of USUV[4] and WNV lineage 2[5]. At the end of the mosquito season, RNA from these samples were pooled and tested for the presence of WNV lineage 1[7].

#### Viral whole genome sequencing and phylogenetic analysis

Consensus sequences were generated using the following approaches. Until end 2019, reference based alignment against an arbitrary chosen Usutu virus genome was performed in Geneious[8] or in CLC Genomic Workbench 11.0 (https://www.qiagenbioinformatics.com/), followed by a second reference based alignment to the most closely related sequence to the consensus sequence identified using Blastn[9,10] and generation of a final consensus sequence. From 2020 reference-based alignment against the most similar reference genome from a selected reference set was performed using Minimap2[11]. A consensus genome was extracted, reads were remapped to the generated consensus sequence, and a new consensus sequence was generated. Positions with a coverage below 100 reads and later below 30 reads were replaced with an ‘N’. Homopolymeric and primer binding regions were manually checked and resolved by consulting raw reads and reference genomes.

All available near full length USUV and WNV lineage 2 genomes were retrieved from GenBank [12] in May 2024. Sequences derived from experimental infections and a set of sequences potentially affected by bioinformatic inaccuracies were excluded. Public sequences were aligned with the newly generated sequences using MUSCLE[13]. A maximum likelihood phylogenetic analysis was performed using IQ-TREE with the substitution models GTR+I+F+G4 for USUV and GTR+F+R4 for WNV lineage 2, with ultrafast bootstrap support calculated using 1,000 replicates[14,15].

##### Antibody detection in serum from live free-ranging birds

In the protein microarray[16], sera were diluted at a 1:80 ratio and tested for IgY reactivity against USUV and WNV NS1 proteins. A signal ≥ 6000 relative fluorescence units (RFU) on the protein microarray was considered above background. Sera with a signal ≥ 30'000 RFU (near half of the maximum fluorescence value, 67000 RFU and corresponding to an approximate titer of 1:80) were considered highly reactive.

In total, 928 sera with a signal above background on the protein microarray were confirmed using neutralization assays. Initially, Virus Neutralization Tests (VNT) were used. Starting in 2020, Focus Reduction Neutralization Tests (FRNT) replaced VNT as primary confirmation method due to increased sensitivity and specificity. To ensure consistency, all samples previously tested on the VNT with sufficient sample volume remaining were retrospectively re-tested using FRNT. For samples tested on the VNT with insufficient material remaining, the VNT result was retained. Consequently, the final serology results are derived from 785 samples tested with FRNT and 143 samples tested with VNT.

Sera were considered positive on the VNT when at least two-serial dilutions showed complete blocking of viral infection (cut-off titer ≥16). Sera were considered positive on the focus reduction neutralization test when titer was ≥1:160 for USUV and ≥1:80 for WNV (calculated based on a ≥70% reduction in infected cells counts).

##### Spatial analysis and cartography

Coordinates of unique events of live birds sampling were clustered spatially using Density-Based Spatial Clustering of Applications with Noise, with parameters set to ε=3000m and minimum points=8. Total number of birds tested were grouped and summarized per cluster. Reports of dead wild birds were aggregated into a hexagonal grid with cells of 30 $km^2$ to group and mask specific locations. Captive dead birds were aggregated per Zoo, and when birds were owned by privates, by location name. Exact coordinates of mosquito traps were recorded, and samples were aggregated by trapping location.

#### Results

##### Avian Host Species with Usutu Virus or West Nile Virus Occurrence

In addition to Common Blackbirds, Song Thrushes (*Turdus philomelos)*, Eurasian Jays (*Garrulus glandarius*), House sparrows (*Passer domesticus*) and Blue Tits (*Cyanistes caeruleus*) showed high USUV prevalence in live birds (0.2% to 1.8%), with infections detected in several dead birds from these species (ranging from 3 to 7). With the exceptions of Blue Tits, USUV neutralizing antibodies were also detected in these species, with seroprevalence ranging from 4.5% to 0.9%. USUV prevalence was 0.6% in live Redwings (*Turdus iliacus*) and 0.3% in live Great Tits (*Parus major*), with no infections detected in dead birds from these species. Reversely, seven Common Wood Pigeons (*Columba palumbus,* order Columbiformes) were found dead with USUV infection, while no infections were detected in the small number of live birds tested (n=61). USUV and WNV detection overlapped in 5 species: Song Thrushes, House Sparrows, Great Tits, Common Whitethroats (*Sylvia communis*) and Common Chiffchaffs (*Phylloscopus collybita*). The complete breakdown of number of individual sampled, viral detections and antibodies detections per bird orders and bird species is presented in **appendix 2.**

**Table S1. Mosquito species sampled and tested at bird ringing sites and via statutory sampling in central Netherlands**

| | Species | N pools tested RT-PCR | N individuals tested RT-PCR | N pools USUV RNA positive | N pools WNV RNA positive | Percent USUV RNA positive pools | Percent WNV RNA positive pools |
|---|---|---|---|---|---|---|---|
| 1 | *Culex pipiens/torrentium* | 4946 | 35089 | 27 | 6 | 0.5% | 0.1% |
| 2 | *Culiseta annulata* | 451 | 1078 | 0 | 0 | 0.0% | 0.0% |
| 3 | *Anopheles claviger* | 233 | 632 | 0 | 0 | 0.0% | 0.0% |
| 4 | *Anopheles maculipennis s.l.* | 181 | 257 | 1 | 0 | 0.6% | 0.0% |
| 5 | *Coquillettidia richiardii* | 135 | 323 | 0 | 0 | 0.0% | 0.0% |
| 6 | *Aedes cinereus* | 109 | 612 | 0 | 0 | 0.0% | 0.0% |
| 7 | *Culiseta morsitans* | 102 | 279 | 0 | 0 | 0.0% | 0.0% |
| 8 | *Culiseta ochroptera* | 98 | 921 | 0 | 0 | 0.0% | 0.0% |
| 9 | *Anopheles plumbeus* | 83 | 146 | 0 | 0 | 0.0% | 0.0% |
| 10 | *Aedes vexans* | 41 | 107 | 0 | 0 | 0.0% | 0.0% |
| 11 | *Culex territans* | 23 | 45 | 0 | 0 | 0.0% | 0.0% |
| 12 | *Culex sp.* | 21 | 111 | 0 | 0 | 0.0% | 0.0% |
| 13 | *Aedes sticticus* | 12 | 20 | 0 | 0 | 0.0% | 0.0% |
| 14 | *Aedes cantans* | 10 | 17 | 0 | 0 | 0.0% | 0.0% |
| 15 | *Aedes rusticus* | 9 | 9 | 0 | 0 | 0.0% | 0.0% |
| 16 | *Aedes annulipes* | 8 | 10 | 0 | 0 | 0.0% | 0.0% |
| 17 | *Aedes sp.* | 5 | 7 | 0 | 0 | 0.0% | 0.0% |
| 18 | *Culex modestus* | 5 | 15 | 0 | 0 | 0.0% | 0.0% |
| 19 | *Anopheles sp.* | 2 | 4 | 0 | 0 | 0.0% | 0.0% |
| 20 | *Culiseta sp.* | 2 | 11 | 0 | 0 | 0.0% | 0.0% |
| 21 | *Aedes communis* | 1 | 1 | 0 | 0 | 0.0% | 0.0% |
| 22 | *Aedes detritus* | 1 | 1 | 0 | 0 | 0.0% | 0.0% |
| 23 | *Aedes punctor* | 1 | 3 | 0 | 0 | 0.0% | 0.0% |
| 24 | *Aedes rossicus* | 1 | 1 | 0 | 0 | 0.0% | 0.0% |
| | **Total** | **6480** | **39699** | **28** | **6** | **0.4%** | **0.1%** |

**Table S2. Number of samples tested and molecular and serological detections of Usutu virus and West Nile virus by research project and year.**

| | 2016 | 2017 | 2018 | 2019 | 2020 | 2021 | 2022 | Total |
|---|---|---|---|---|---|---|---|---|
| **Live birds - landbirds** | | | | | | | | |
| N tested USUV and WNV PCR | 627 | 975 | 1574 | 2203 | 5022 | 6151 | 6148 | 22700 |
| N USUV RNA positive | 12 | 21 | 39 | 22 | 5 | 2 | 55 | 156 |
| N WNV RNA positive | 0 | 0 | 0 | 0 | 8 | 0 | 0 | 8 |
| Percent USUV RNA positive | 1.9% | 2.2% | 2.5% | 1.0% | 0.1% | 0.0% | 0.9% | 0.7% |
| Percent WNV RNA positive | 0.0% | 0.0% | 0.0% | 0.0% | 0.2% | 0.0% | 0.0% | 0.0% |
| N tested USUV and WNV PMA | 94 | 320 | 464 | 417 | 657 | 786 | 1438 | 4176 |
| N USUV antibodies positive | 3 | 35 | 59 | 51 | 20 | 31 | 41 | 240 |
| N WNV antibodies positive | 1 | 6 | 9 | 2 | 18 | 8 | 9 | 53 |
| N *Orthoflavivirus*-neutralizing antibodies positive | 0 | 5 | 3 | 4 | 15 | 5 | 8 | 40 |
| Percent USUV-neutralizing antibodies positive* | 3.2% | 10.9% | 12.7% | 12.2% | 3.0% | 3.9% | 2.9% | 5.7% |
| Percent WNV-neutralizing antibodies positive* | 1.1% | 1.9% | 1.9% | 0.5% | 2.7% | 1.0% | 0.6% | 1.3% |
| Percent *Orthoflavivirus*-neutralizing antibodies antibodies positive* | 0.0% | 1.6% | 0.6% | 1.0% | 2.3% | 0.6% | 0.6% | 1.0% |
| **Live birds - waterbirds** | | | | | | | | |
| N tested USUV and WNV PCR | 27 | 38 | 192 | 286 | 618 | 3577 | 5980 | 10718 |
| N USUV RNA positive | 0 | 0 | 1 | 0 | 0 | 1 | 1 | 3 |
| N WNV RNA positive | 0 | 0 | 0 | 0 | 0 | 0 | 1 | 1 |
| Percent USUV RNA positive | 0.0% | 0.0% | 0.5% | 0.0% | 0.0% | 0.0% | 0.0% | 0.0% |
| Percent WNV RNA positive | 0.0% | 0.0% | 0.0% | 0.0% | 0.0% | 0.0% | 0.0% | 0.0% |
| N tested USUV and WNV PMA | | | | 19 | 71 | 27 | 26 | 143 |
| N USUV antibodies positive | | | | 0 | 0 | 1 | 2 | 3 |
| N WNV antibodies positive | | | | 0 | 0 | 0 | 0 | 0 |
| N *Orthoflavivirus*-neutralizing antibodies positive | | | | 0 | 2 | 0 | 2 | 4 |
| Percent USUV-neutralizing antibodies positive* | | | | 0.0% | 0.0% | 3.7% | 7.7% | 2.1% |
| Percent WNV-neutralizing antibodies positive* | | | | 0.0% | 0.0% | 0.0% | 0.0% | 0.0% |
| Percent *Orthoflavivirus*-neutralizing antibodies antibodies positive* | | | | 0.0% | 2.8% | 0.0% | 7.7% | 2.8% |
| **Dead birds - free-ranging** | | | | | | | | |
| N tested USUV PCR | 94 | 110 | 119 | 95 | 230 | 243 | 289 | 1180 |
| N tested WNV PCR | 5 | 18 | 79 | 89 | 230 | 243 | 289 | 953 |
| N USUV RNA positive | 53 | 38 | 72 | 8 | 15 | 3 | 54 | 243 |
| Percent USUV RNA positive | 56.4% | 34.5% | 60.5% | 8.4% | 6.5% | 1.2% | 18.7% | 20.6% |
| **Dead birds - captive** | | | | | | | | |
| N tested USUV PCR | 19 | 10 | 28 | 112 | 175 | 208 | 101 | 653 |
| N tested WNV PCR | 0 | 1 | 23 | 112 | 175 | 208 | 101 | 620 |
| N USUV RNA positive | 13 | 6 | 7 | 5 | 2 | 4 | 4 | 41 |
| Percent USUV RNA positive | 68.4% | 60.0% | 25.0% | 4.5% | 1.1% | 1.9% | 4.0% | 6.3% |
| **Mosquitoes - Central Netherlands** | | | | | | | | |
| N pools tested USUV and WNV PCR | | | | 619 | 1824 | 1349 | 998 | 4790 |
| N individuals tested USUV and WNV PCR | | | | 3422 | 8922 | 8842 | 6033 | 27219 |
| N pools USUV RNA positive | | | | 5 | 0 | 1 | 12 | 18 |
| N pools WNV RNA positive | | | | 0 | 1 | 0 | 0 | 1 |
| Percent pools USUV RNA positive | | | | 0.8% | 0.0% | 0.1% | 1.2% | 0.4% |
| Percent pools WNV RNA positive | | | | 0.0% | 0.1% | 0.0% | 0.0% | 0.0% |
| **Mosquitoes - Bird research stations** | | | | | | | | |
| N pools tested USUV and WNV PCR | | | | | 415 | 809 | 466 | 1690 |
| N individuals tested USUV and WNV PCR | | | | | 3488 | 5621 | 3371 | 12480 |
| N pools USUV RNA positive | | | | | 1 | 0 | 9 | 10 |
| N pools WNV RNA positive | | | | | 5 | 0 | 0 | 5 |
| Percent pools USUV RNA positive | | | | | 0.2% | 0.0% | 1.9% | 0.6% |
| Percent pools WNV RNA positive | | | | | 1.2% | 0.0% | 0.0% | 0.3% |

PMA = protein microarray
*Percentage of samples tested on the protein microarray that were confirmed positive for neutralizing antibodies by focus reduction neutralization test or virus neutralization test

**Figure S1. Geographical distribution of Usutu virus (USUV) detections in birds in the Netherlands per year, 2016-2022**

Square symbols indicate detections in dead captive birds, and circular symbols detections in live birds, with sizes proportional to the number of detections. Shading intensity in hexagonal grid cells (size: 30 km²) reflect the number of detections in dead free-ranging birds in that cell. Source administrative boundaries: CBS, Kadaster, "CBSGebiedsindelingen2022" (https://service.pdok.nl/cbs/gebiedsindelingen/atom/v1_0/index.xml).

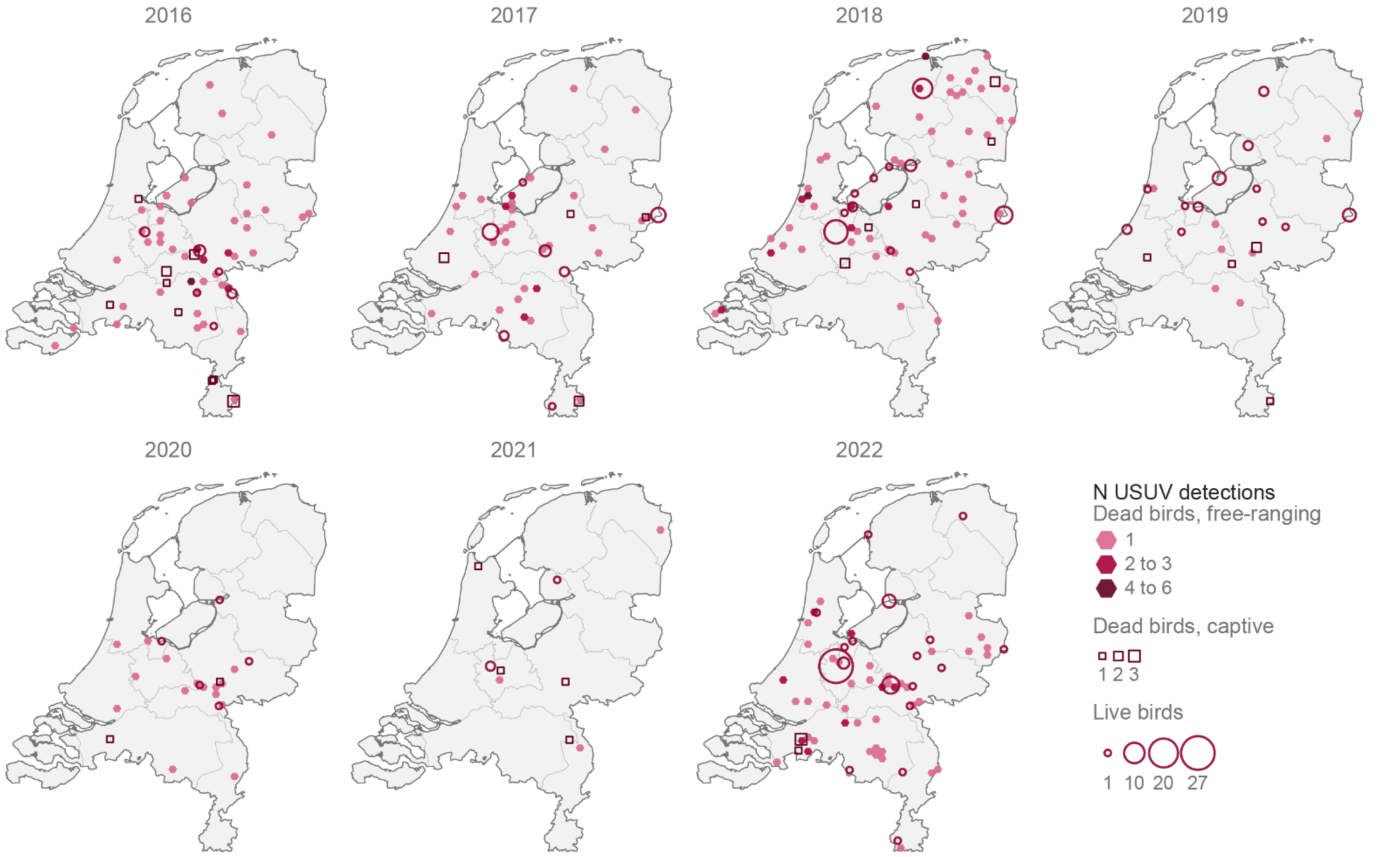

**Figure S2. Geographical distribution of Usutu virus (USUV), West Nile virus (WNV) and *Orthoflaviviruses*-neutralizing antibodies detections in live free-ranging birds per year in the Netherlands, 2016-2022**

Circular symbols are coloured per type of antibody and sized proportionally to the number of detections. Source administrative boundaries: CBS, Kadaster, "CBSGebiedsindelingen2022" (https://service.pdok.nl/cbs/gebiedsindelingen/atom/v1_0/index.xml).

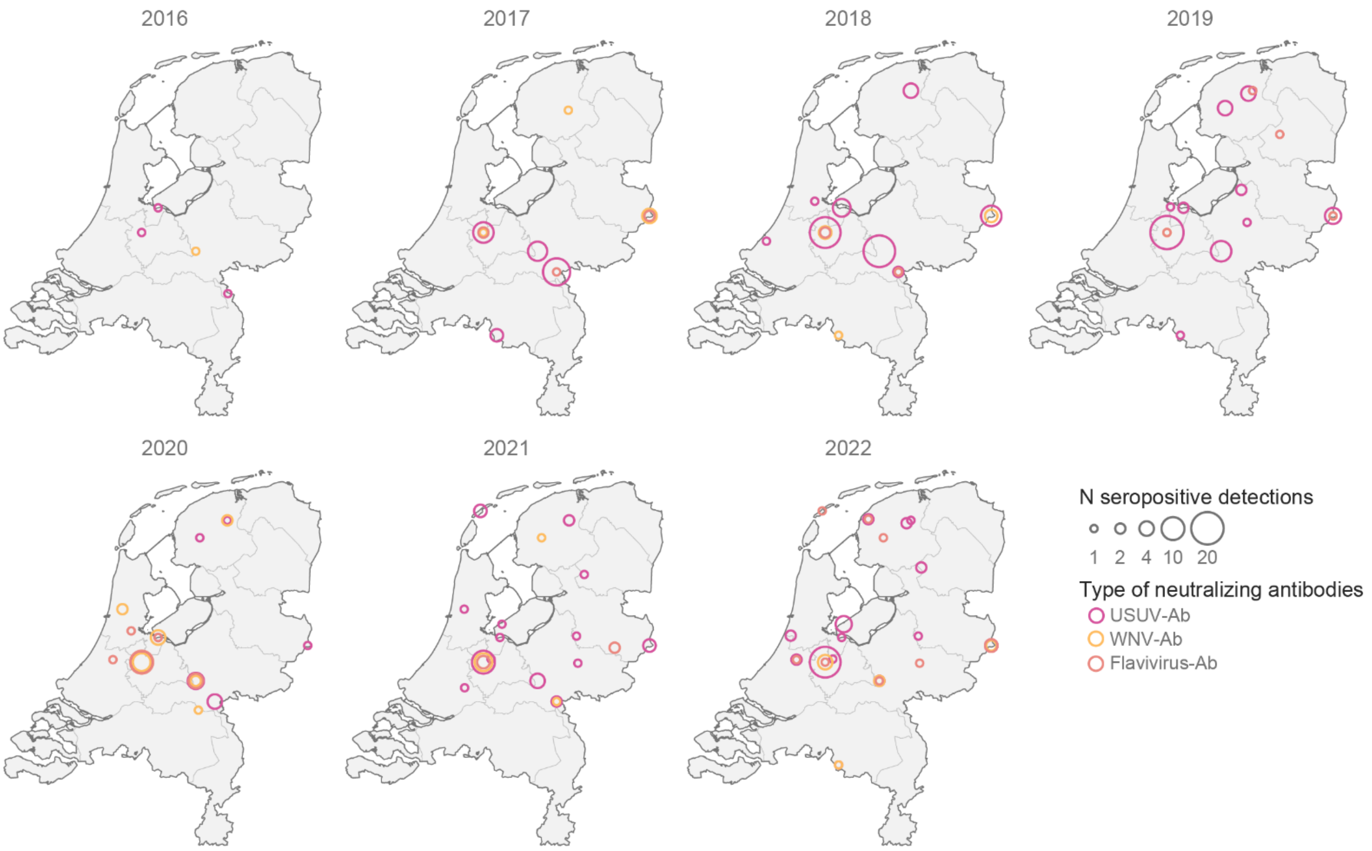

**Figure S3. Annual distribution of live bird serum sampling and Usutu virus-neutralizing antibody (USUV-neutralizing Ab) seroprevalence by bird ringing area, the Netherlands, 2016-2022**

Circular symbols indicate sampling areas from where live bird serum samples were tested for *Orthoflaviviruses* antibodies on the protein microarray, with symbol size corresponding to the number of samples tested. Color intensity indicates the percentage of these samples confirmed positive for USUV-neutralizing Ab by focus reduction neutralization test or virus neutralization test. Source administrative boundaries: CBS, Kadaster, "CBSGebiedsindelingen2022" (https://service.pdok.nl/cbs/gebiedsindelingen/atom/v1_0/index.xml).

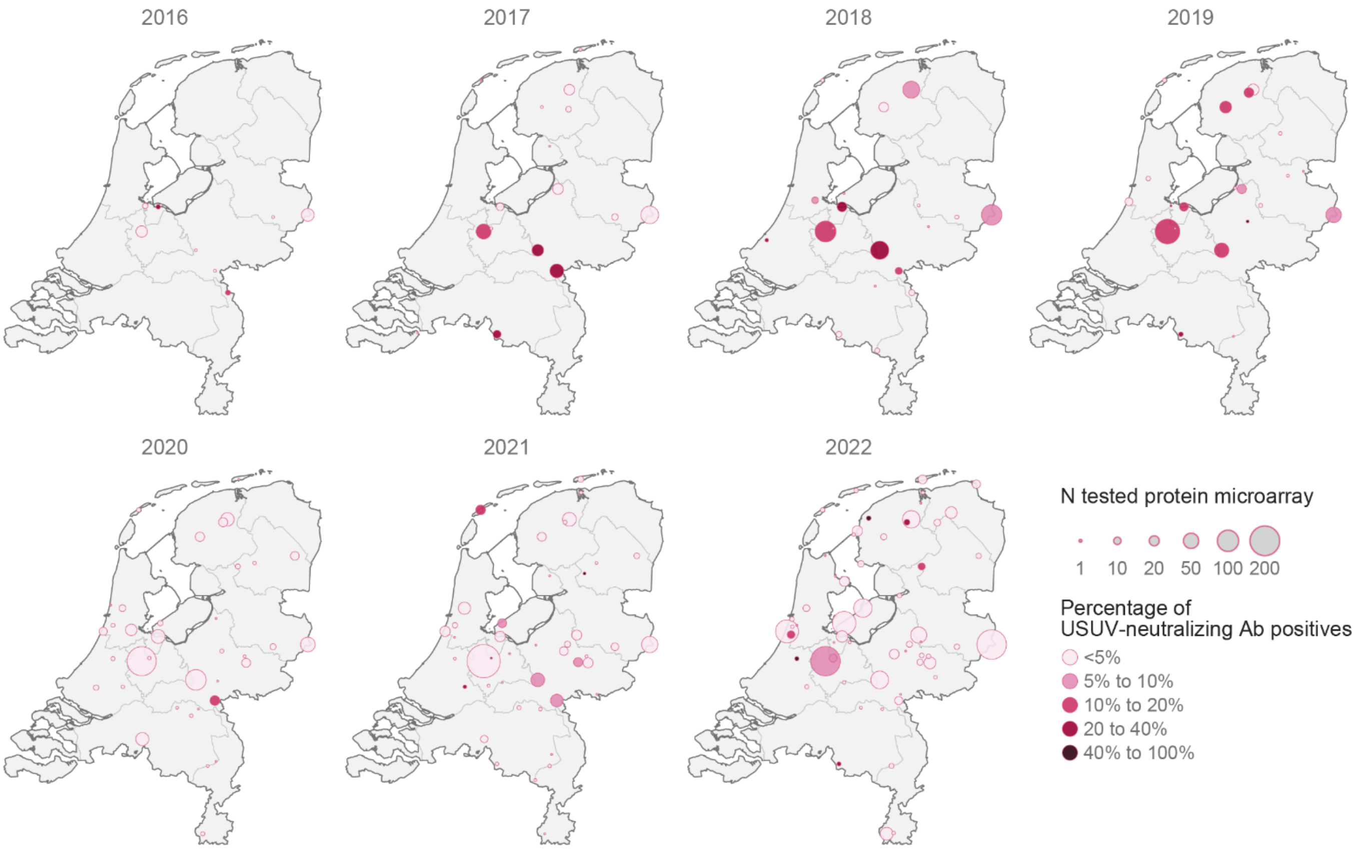

**Figure S4. Geographical distribution of West Nile virus (WNV) and neutralizing antibodies (WNV-neutralizing Ab) detections in live birds and mosquitoes in the Netherlands per year, 2016-2022**

Orange symbols indicate molecular detections in live birds (solid circles, sized proportionally to the number of detections) and mosquitoes (cross). Circular outline symbols represent WNV-Ab detections (sized proportionally to the number of detections), with yellow indicating all detections and purple highlighting detections in local birds. Source administrative boundaries: CBS, Kadaster, "CBSGebiedsindelingen2022" (https://service.pdok.nl/cbs/gebiedsindelingen/atom/v1_0/index.xml).

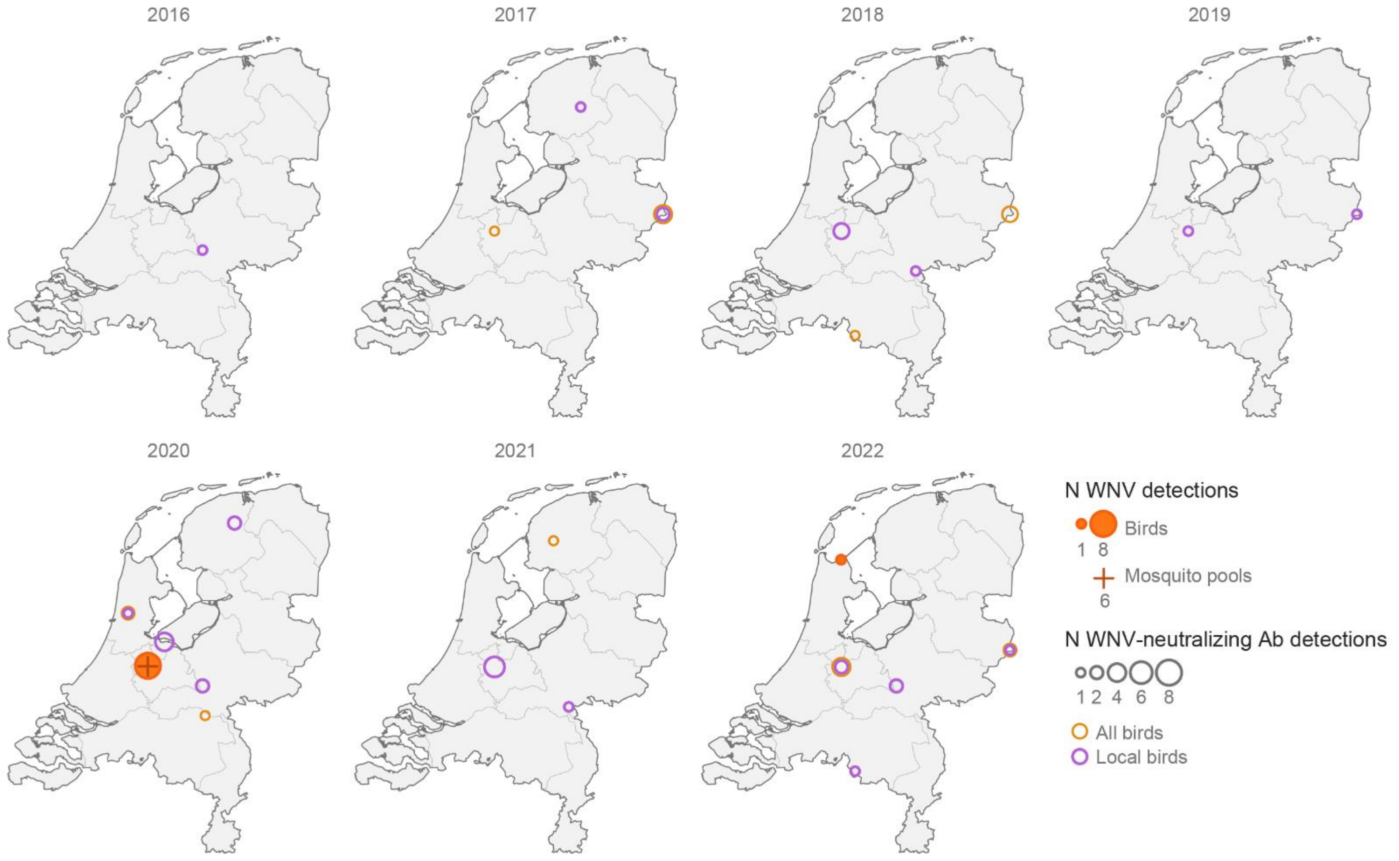

**Figure S5. Geographical distribution of live free-ranging birds captures sampled and tested per year in the Netherlands, 2016-2022**

Circular symbols are sized proportionally to the number of birds tested per sampling area. The color scheme indicates the predominant bird types sampled. Source administrative boundaries: CBS, Kadaster, "CBSGebiedsindelingen2022" (https://service.pdok.nl/cbs/gebiedsindelingen/atom/v1_0/index.xml).

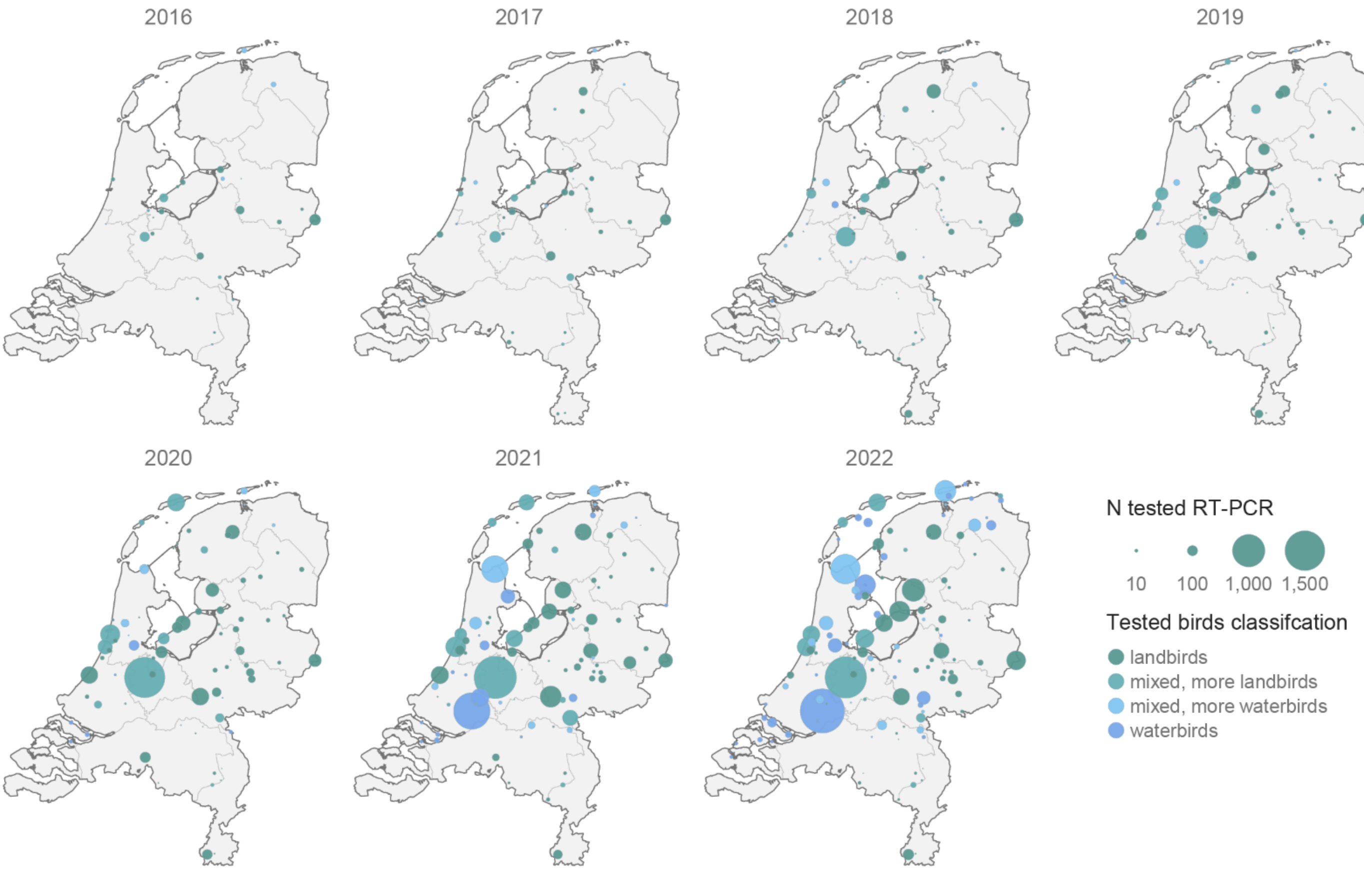

**Figure S6. Geographical distribution of dead free-ranging birds and dead captive birds tested per year in the Netherlands, 2016-2022**

Square symbols indicate sites of sampling of dead captive birds, with sizes proportional to the number of birds tested. Shading intensity in hexagonal grid cells (size: 30 km²) reflect the number dead free-ranging birds sampled in that cell. Source administrative boundaries: CBS, Kadaster, "CBSGebiedsindelingen2022" (https://service.pdok.nl/cbs/gebiedsindelingen/atom/v1_0/index.xml).

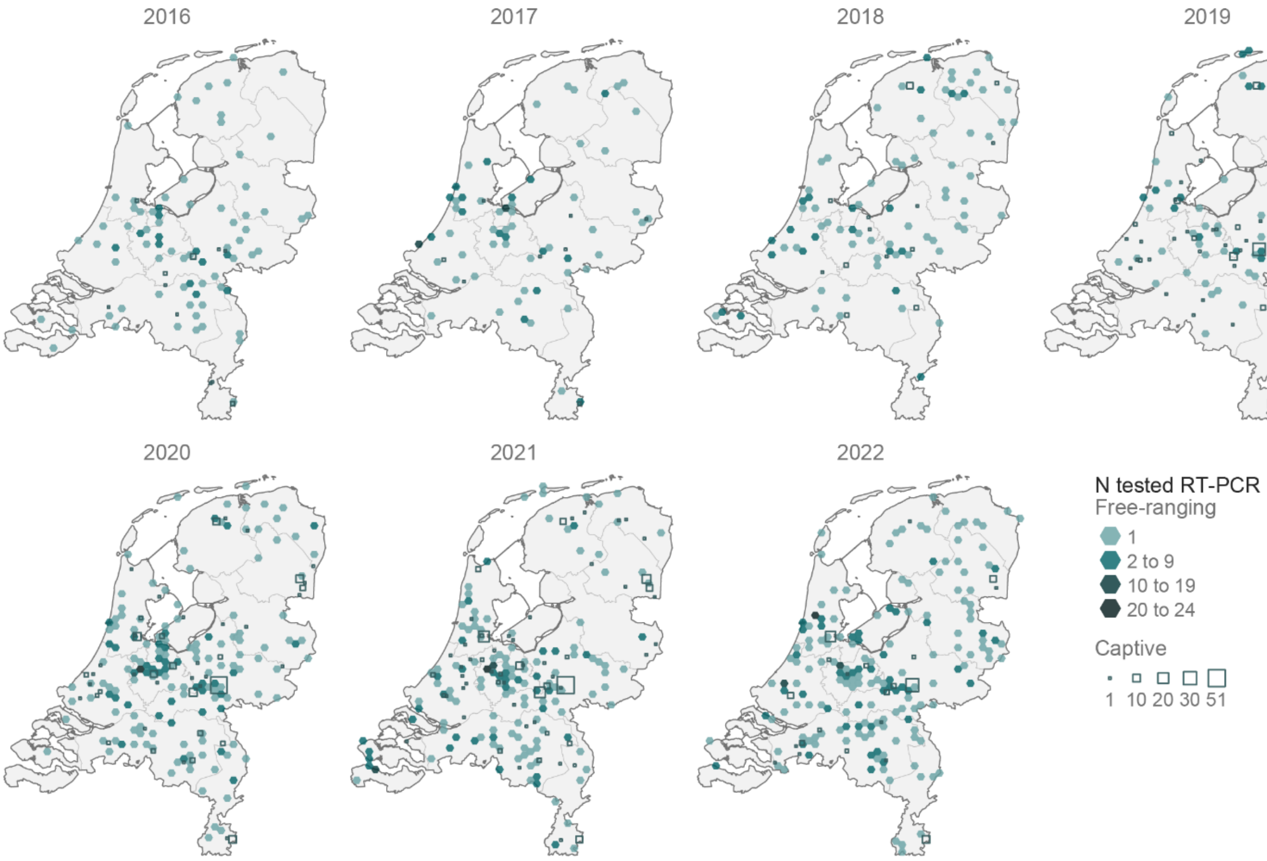
